# Identification and characterization of a potentially novel dorsal cutaneous muscle in rodents

**DOI:** 10.1101/2024.01.30.577894

**Authors:** Jeffrey C. Petruska

## Abstract

In the course of performing a detailed dissection of adult rat to map the cutaneous nerves of cervical, thoracic, and lumbar levels a small and unexpected structure was isolated. It appeared to be a cutaneous striated muscle and was observed in both male and female rats and in mice but absent from cats and humans. With the skin reflected laterally from midline, the muscle lies closely apposed to the lateral border of the Thoracic Trapezius (Spinotrapezius) muscle and is easily missed in standard gross dissections. Focussed prosections were performed to identify the origin, insertion, and course of gross innervation. Identification of each of these elements showed them to be distinct from the nearby Trapezius and Cutaneous Trunci (Cutaneous Maximus in mouse) muscles. The striated muscle nature of the structure was validated with whole-mount microscopy.Consulting a range of published rodent anatomical atlases and gross anatomical experts revealed no prior descriptions. This preliminary report is an opportunity for the anatomical and research communities to provide input to either confirm the novelty of this muscle or refer to prior published descriptions in rodents or other species while the muscle, its innervation, and function are further characterized. Presuming this muscle is indeed novel, the name “Cutaneous Scapularis muscle” is proposed in accord with general principles of the anatomical field.

## Introduction

Detailed dissections were carried out for the purpose of creating an in-house pictorial atlas of cutaneous nerves and the landmarks that could be used to identify them for surgical and tissue-retrieval purposes. In the course of these dissections an unexpected small cutaneous muscle was encountered.

For the purposes here I will assume the muscle is novel and has not previously been assigned a name. Although anatomists do not agree on all details, there are some common principles that one can follow to suggest useful names for previously un-named structures and structures with multiple names (e.g., [1]). The proposed name – Cutaneous scapularis muscle – reflects both the nature and position of the muscle in the rat (species of original identification) and uses a singularly-named reference point (scapula).

## Methods

### Animals underwent prosection to enable documentation

All animals involved were included in approved IACUC protocols. Adult male and female Sprague-Dawley rats were euthanized with CO_2_ and prosections were performed for characterization and to identify the origin, insertion, and innervation of the Cutaneous Scapularis muscle (n=2). Confirmatory dissections were performed with additional adult male and female Sprague-Dawley rats (n>10), including a Thy1-GFP rat to gain a perspective of the native innervation. Comparative dissections were performed to determine if the Cutaneous Scapularis (or equivalent) might also exist in cats (n=2) and mice (n>6), including Thy1-YFP mice. Images were captured with digital cameras (handheld or mounted to surgical microscope or inverted fluorescence microscope) and modified only for cropping/orientation and for brightness, contrast, and color balance.

### Resources were consulted to determine novelty of the finding

In order to determine if the muscle had been identified and characterized previously, a variety of resources were consulted. Paper and online text and image-based compendia and databases were consulted including Nomina Anatomica Veterinaria (6^th^ edition) [2], IMAIOS [3], Google, and anatomical atlases [4, 5].

Details of the muscle were also presented to a conference of professional anatomical educators and individual experts in both human and veterinary anatomy.

## Results

### The Cutaneus scapularis appears to be novel

The text and database resources consulted offered no indication of prior identification or description of the structure-in-question.

Each of the potential cutaneous muscles listed in Nomina Anatomica Veterinaria was assessed for the description of its position and through examination of images retrieved from internet searches of those names. In addition, a general internet search was performed for combinations of rodent/rat/mouse AND cutaneous muscle. A PubMed searched using the terms (novel AND muscle AND gross AND anatomy) yielded >200 results. None of these provided any indication that the muscle had been previously identified. A major limitation of this route of searching is that it was limited to English language.

Details of the muscle were also presented to colleagues with a significant breadth and depth of knowledge. Muscle characteristics were presented as part of one of the keynote seminars at the 2019 joint meeting of the Human Anatomy and Physiology Society (HAPS) and the American Association of Clinical Anatomists (AACA) at Bellarmine University in Louisville, KY. None of the attendees indicated knowledge of the muscle or a suitable homolog or analog in another species.

Elements of these data, including images, were reviewed by a close colleague with extensive knowledge of comparative veterinary anatomy, especially equine anatomy which is highly relevant as horses have some of the most extensive cutaneous musculature on this region [6]. Numerous cutaneous muscles from a range of species were considered but none matched the Cutaneus scapularis.

### Exposure of the Cutaneus scapularis requires specific prosection procedures

Exposure in both rats and mice began with a lateral reflection of the skin from a midline incision. In mice, the muscle is quite small and can be difficult to observe even with aid of a surgical microscope and intent to identify. The Cutaneous scapularis muscle is narrow and long, predominantly tubular, spanning a length of more than 12 vertebrae (approximate range of C5-T10). The following description corresponds to the structure and procedures in rat. The rostral portion of the muscle is often visible without dissection but much of its extent is not visible (Figure 1), and the muscle is easily missed or dismissed as part of the Spinotrapezius. Much of the Cutaneus scapularis muscle is deep to the subcutaneous fat pad, making it necessary to carefully separate the fat pad and remove it from the muscle in order to observe the entire course. Much of its length lies lateral to the lateral border of the Spinotrapezius muscle. The Cutaneus scapularis is more closely apposed to the lateral edge of Spinotrapezius at their mutual caudal extent and becomes increasingly separated from the Spinotrapezius as both muscles extend more rostral (Figure 2). By gross anatomical inspection, the insertion into the dermis appears to overlie the scapula, which was confirmed by section and manipulation of the muscle. The insertion area appeared to be oblong, with a long-axis parallel to the spine (Figure 3). The insertion does not reach midline, unlike that of the nearby Cutaneus trunci muscle (Figure 4). The origin of the Cutaneus scapularis muscle appears to be continuous with the aponeurosis of the caudal Spinotrapezius and was not readily separable by basic dissection (Figure 5).

**Figure 1.**
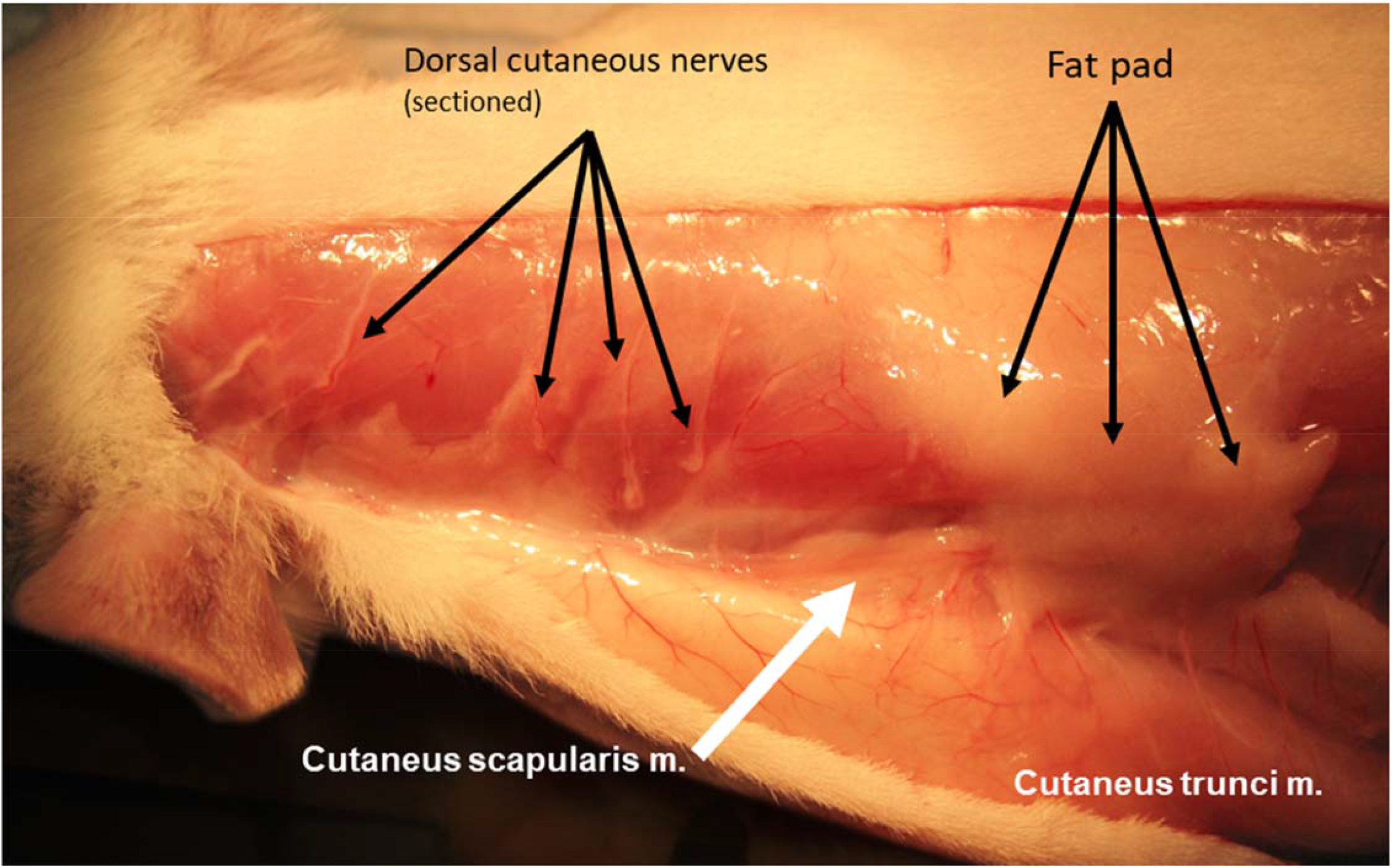
Cutaneus scapularis and Cutaneus trunci muscles and subcutaneous fat pad visualized by reflection of the skin without dissection in rat. Some of the cervical dorsal cutaneous nerves have been sectioned to allow to skin to be reflected laterally without restriction. Thoracic dorsal cutaneous nerves were left intact and can be seen piercing the Cutaneus trunci muscle.

**Figure 2.**
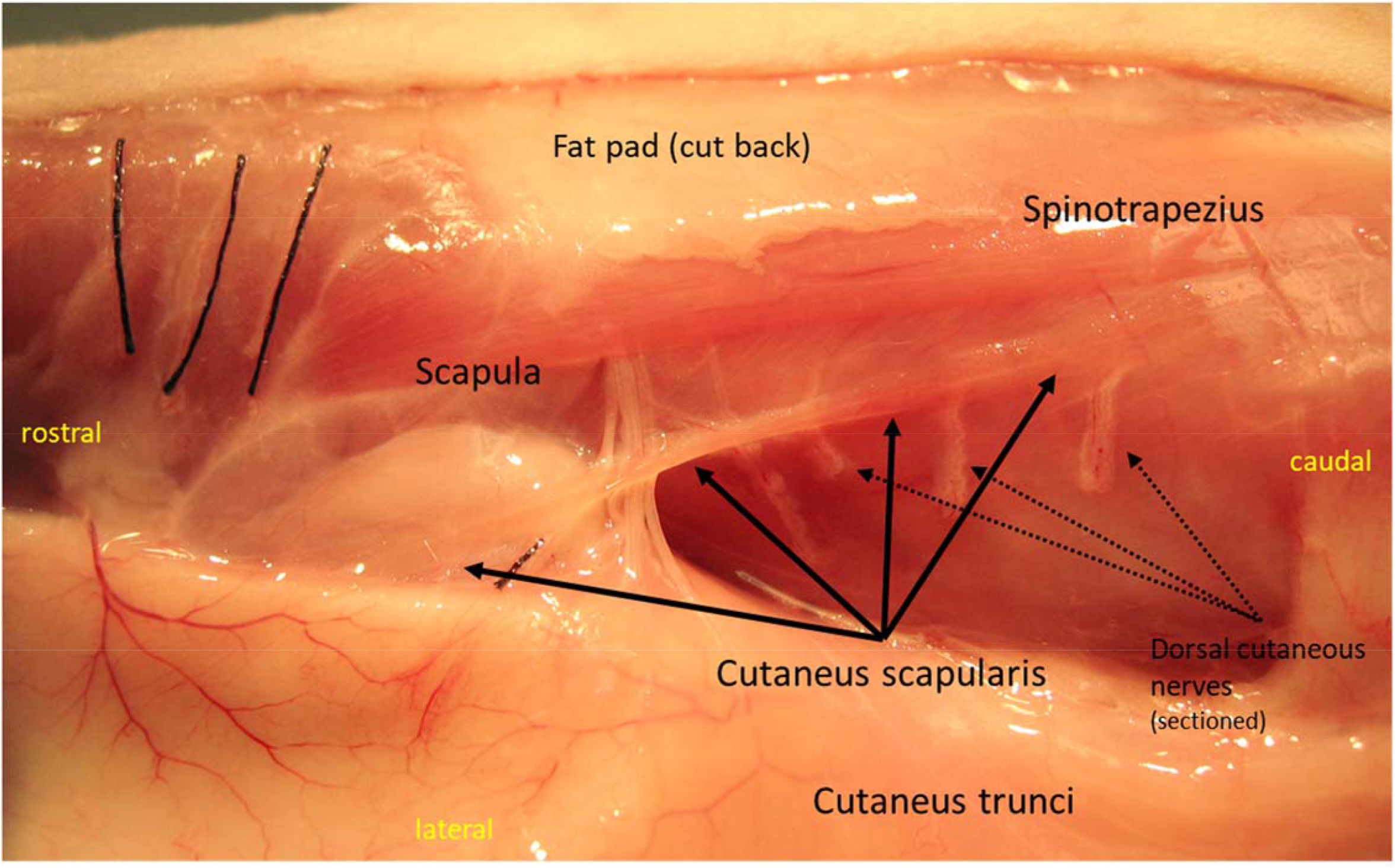
Skin reflection with removal of the overlying dorsal thoracic fat pad reveals the course of the Cutaneus scapularis m.

**Figure 3.**
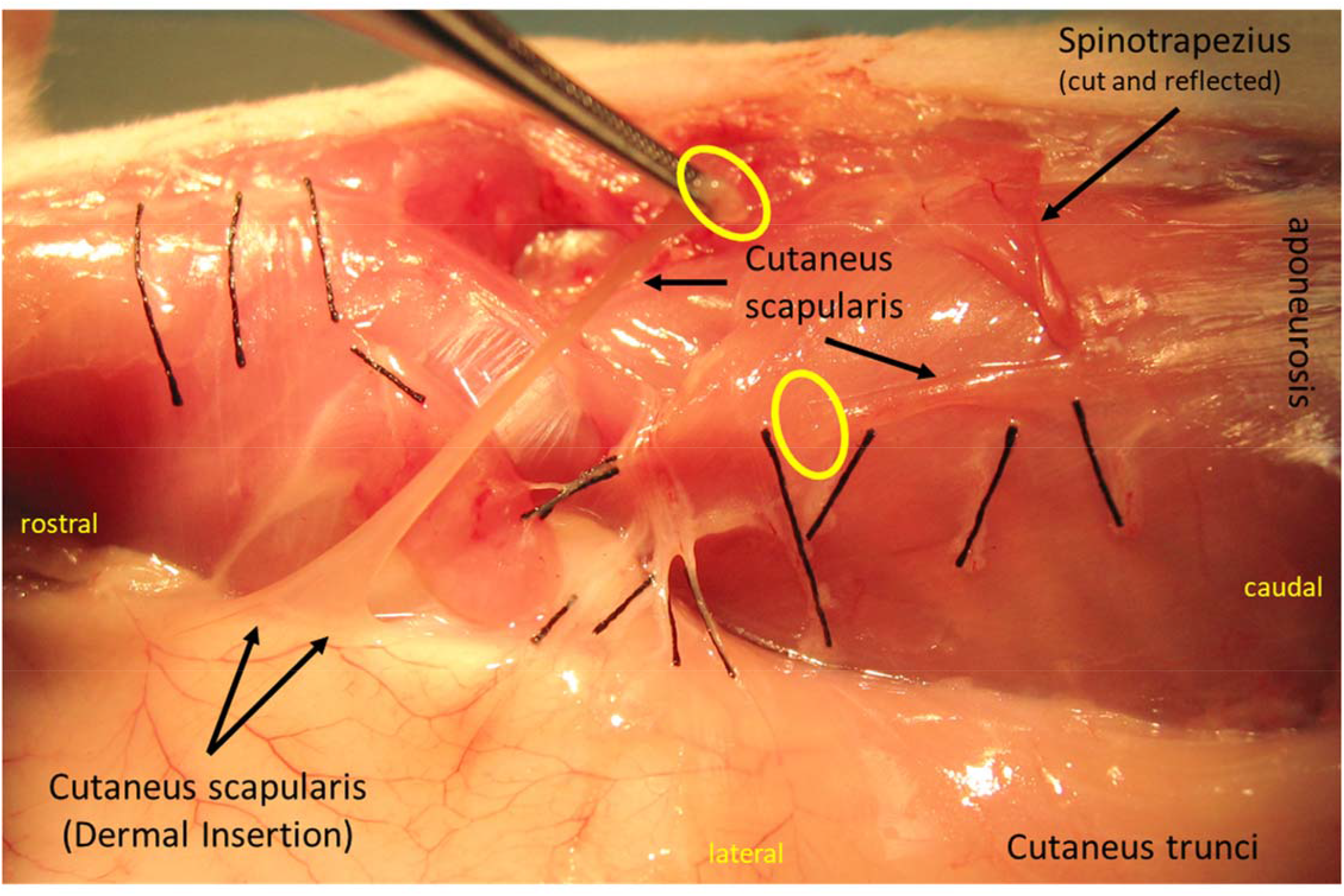
Insertion of the Cutaneus scapularis muscle, which has been cut to enable manipulation to identify connection with the dermis. Yellow ovals indicate the cut ends of the Cutaneus scapularis. Black threads overlie cutaneous nerves.

**Figure 4.**
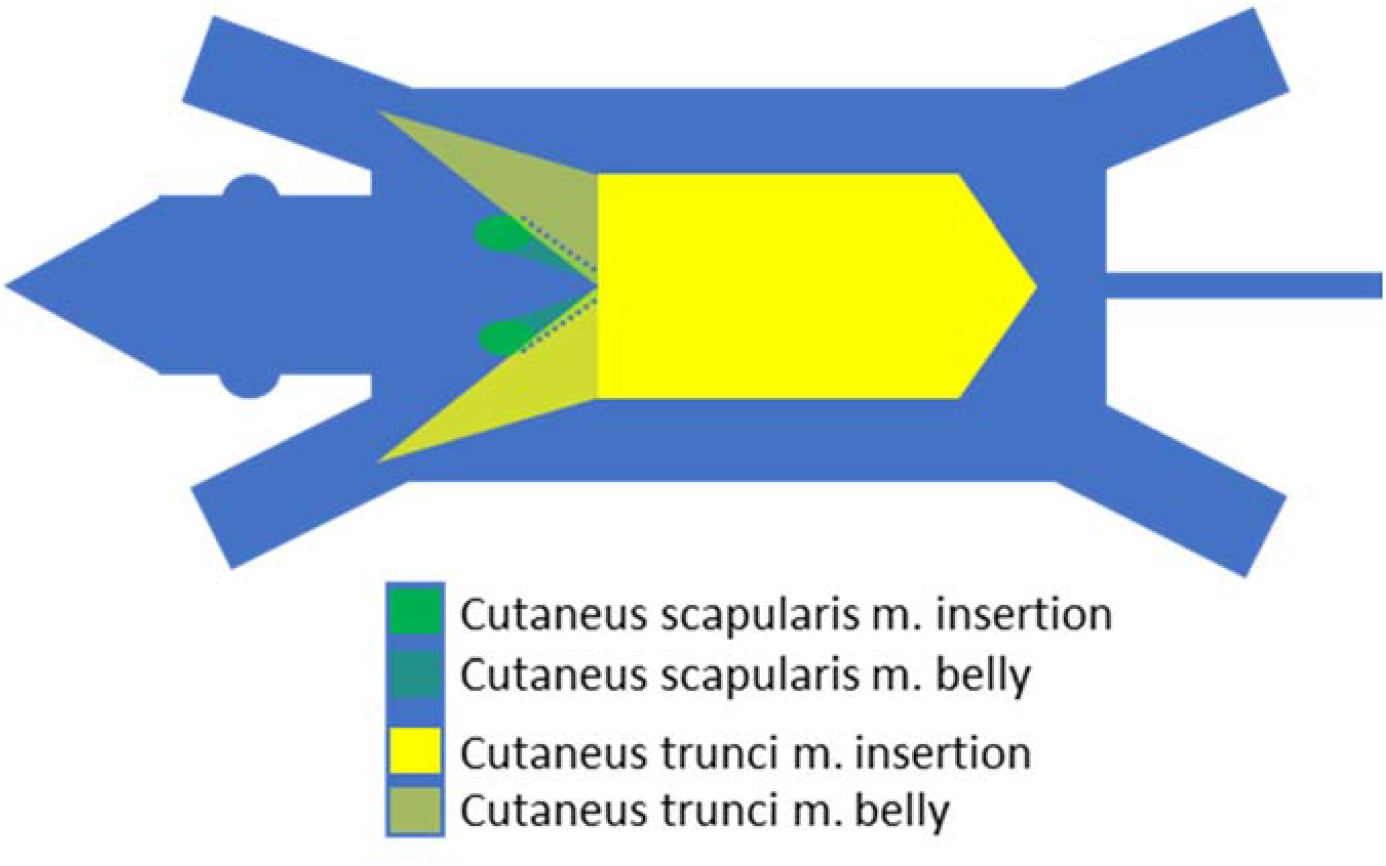
Schematic demonstrating the relationship of the Cutaneus trunci and Cutaneus scapularis muscles.

**Figure 5.**
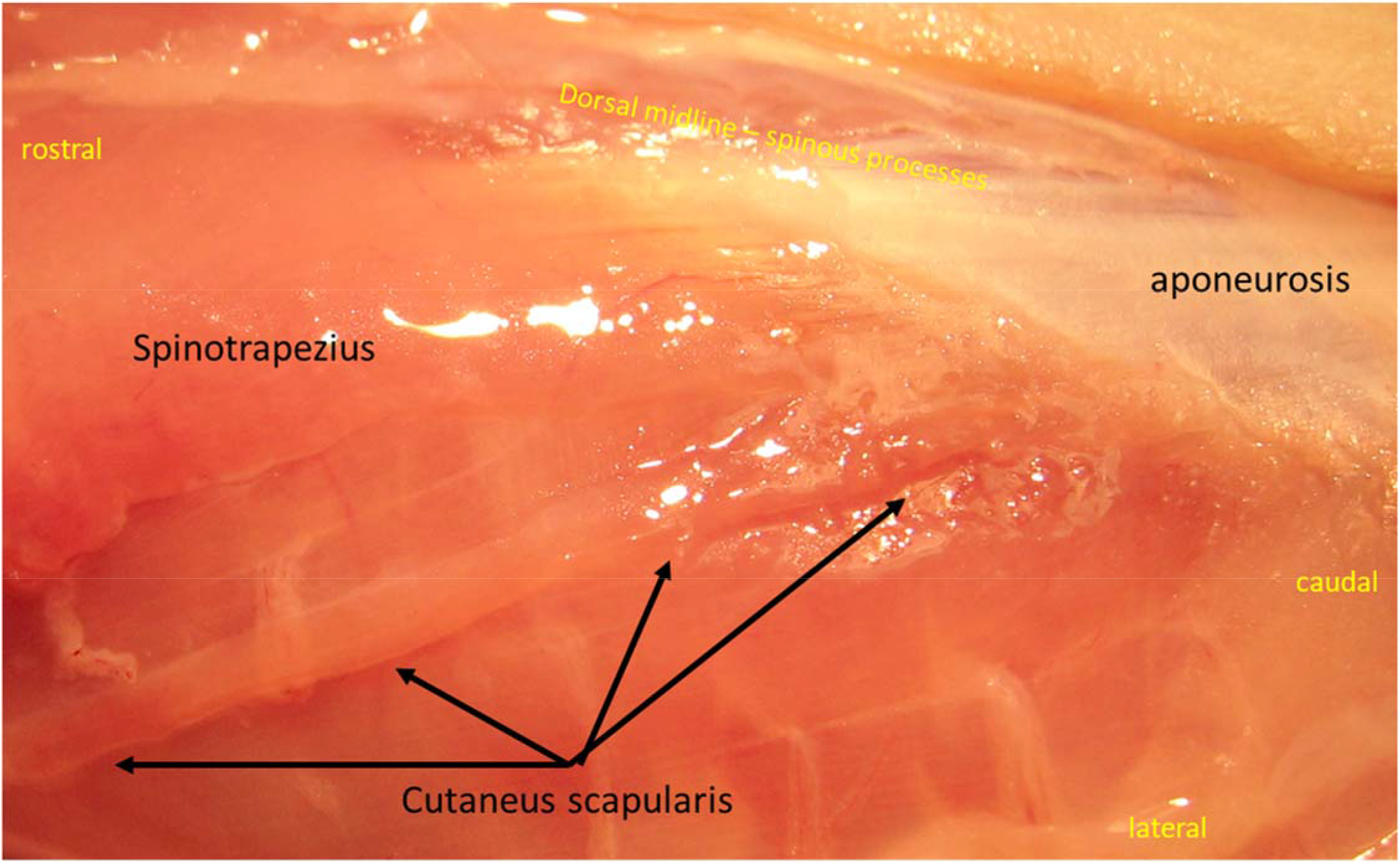
Origin of the Cutaneus scapularis and Spinotrapezius muscles in rat.

The Cutaneous scapularis muscle was sectioned to enable more thorough gross isolation and manipulation to better identify origin, insertion, and any vascular and/or neural attachments (Figure 3). Manipulation revealed a firm mechanical fibrous connection (i.e., more than subcutaneous fascia) between the belly of the Cutaneus scapularis muscle and the body wall. The connection appeared to be a branch of the T4 dorsal cutaneous nerve (Figure 6). Manipulation revealed that this branch maintained a strong connection to the muscle despite the muscle being easily separated from the underlying T1-4 dorsal cutaneous nerves.

**Figure 6.**
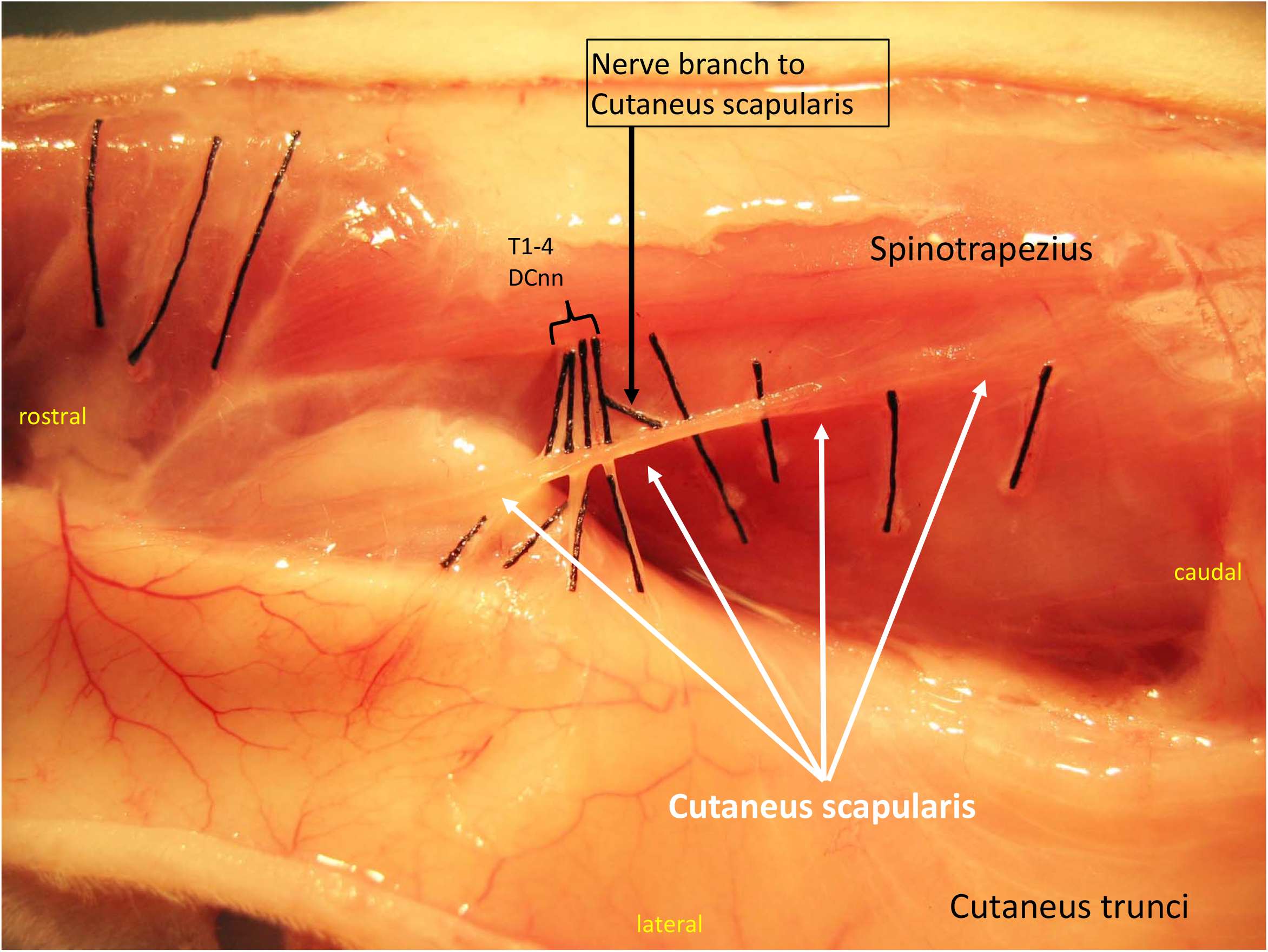
Apparent innervation of the Cutaneus scapularis muscle. This can be seen without markers in Figure 2. Black threads overlie cutaneous nerves. T1-4 DCnn = Dorsal cutaneous nerves of Thoracic segments 1 through 4.

### The Cutaneus scapularis muscle is present in mice

Dissection in mice revealed a structurally and positionally similar muscle (Figure 7). Analogy and homology remain to be determined. The muscle in mice was very small and easily overlooked and damaged. Its apparent insertion into the dermis appeared further rostral and lateral – closer to the base of the ear – than in the rat. Microscopic examination of the isolated muscle revealed separate myofibers and clear striations (Figure 8), confirming that the tissue is indeed skeletal muscle tissue and not smooth muscle or non-muscle tissue.

**Figure 7.**
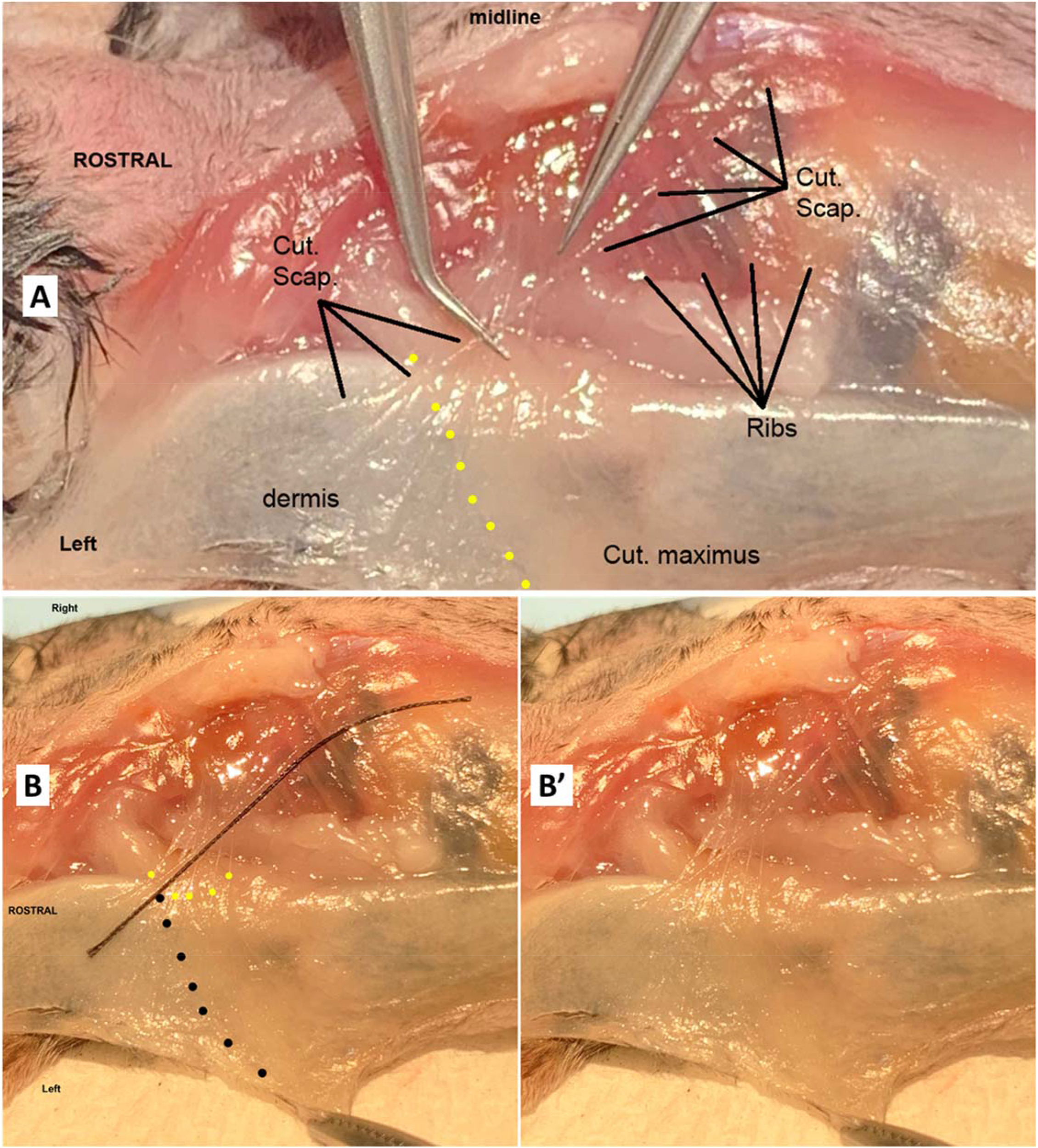
(A) Cutaneus scapularis muscle in adult mouse lifted lightly by forceps. (B) Black thread lies next to Cutaneus scapularis muscle. Black dots indicate the rostral edge of the Cutaneus maximus muscle. (B’) Same field as B but without markup.

**Figure 8.**
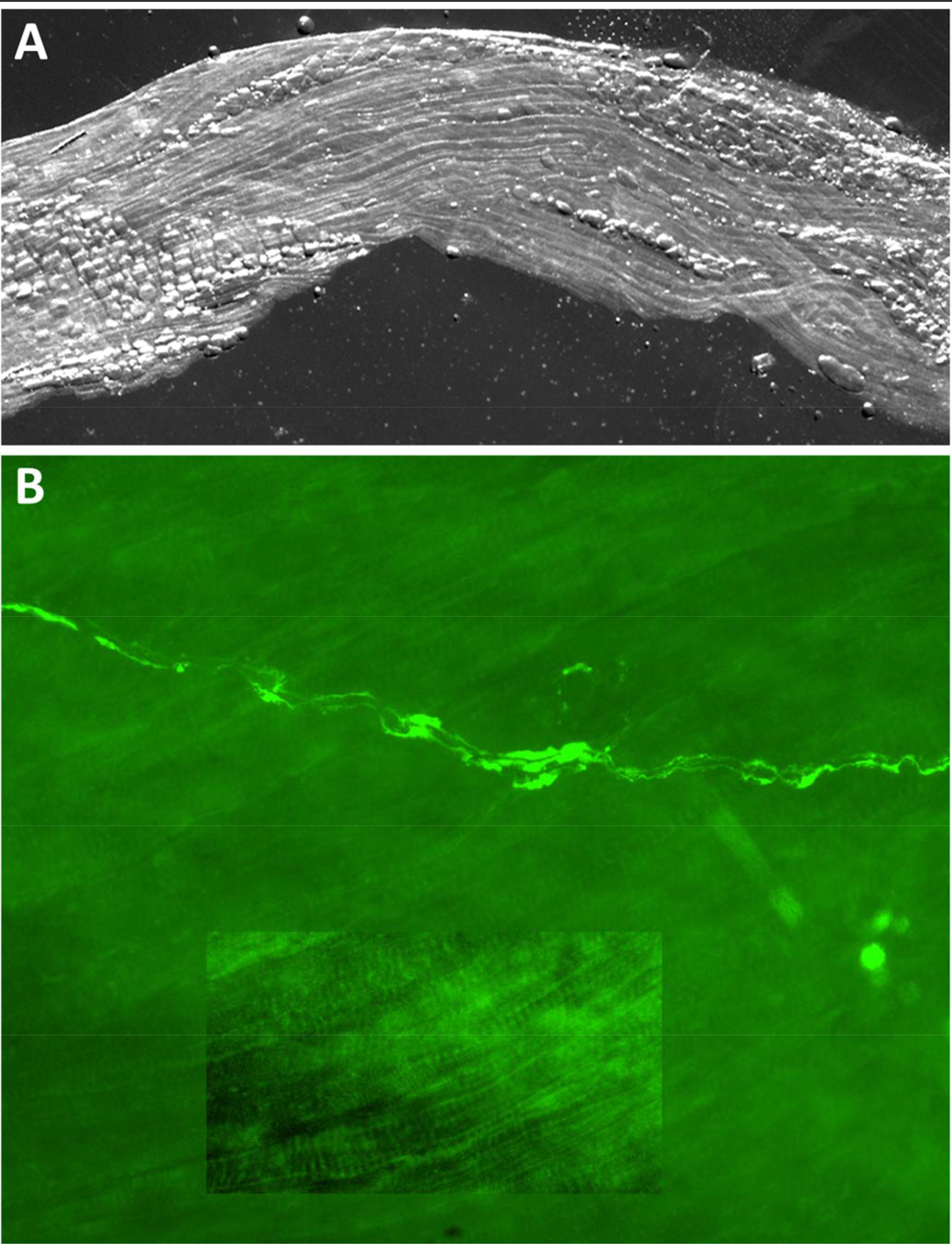
Ex vivo Cutaneus scapularis muscle from adult Thy1-YFP mouse (A) viewed with polarized light and (B) fluorescence filters to reveal YFP. Inset was digitally-enhanced to increase contrast, revealing striations in myofibers.

### The Cutaneus scapularis muscle is not present in cats

Dissection in cats (n=2) revealed no analogous muscle. The expected insertion region and its surround was instead occupied only by the well-documented Platysma muscle, which covers an extensive area of the neck and upper trunk.

## Discussion

### Novelty

It is rare, but not impossible, to make truly novel gross anatomical identifications in highly-studied species in the modern era. For example, a novel peripheral ganglion was identified in mouse with the aid of genetic reporter molecules [7]. It is entirely possible that this muscle has previously been described outside of my knowledge and that of colleagues but has not been included in the consulted atlases or lists of anatomical structures. Even human gross anatomical structures can be omitted from seemingly-encompassing documents [8, 9]. A range of cutaneous muscles have been described and sometimes referred to as the Panniculus carnosus [6]. Panniculus carnosus has been proposed as an umbrella term for cutaneous striated muscles throughout the human body [6], but in a range of non-human species classically refers to muscles that correspond largely to the rat and mouse Cutaneus trunci and Cutaneus maximus, respectively. I examined the range of Paniculus carnosus-like cutaneous muscles reviewed by Naldaiz-Gastes and colleagues [6] and none appeared similar to the Cutaneus scapularis. The description herein clearly indicates that the Cutaneous scapularis is distinct from the rat Cutaneus trunci and mouse Cutaneus maximus [10, 11].

It was also considered that the Cutaneus scapularis muscle might be a homolog or analog of Langer’s axillary arch in human, which has a wide range of characteristics [12, 13]. Although some characteristics appeared similar, the Cutaneus scapularis is too distant from the axilla, and clearly dorsally situated, to be considered a relation to the axillary arch [12, 13].

The Sato and Yaginuma group and colleagues performed an impressive analysis of mammalian embryological PNS development and re-analyses of prior data to propose a new organizational model of nerve development and neuromuscular targeting [14, 15]. Unfortunately, most of the imaging data which might be used to inform the innervation of the Cutaneus scapularis muscle is absent, because the skin was removed. Nonetheless, it appears that the Cutaneus scapularis may be unique in its (eventual) muscular classification and/or its innervation according to the traditional or novel systems. If it is borne out that innervation of the Cutaneus scapularis muscle is via a branch of the T4 dorsal cutaneous nerve, it could imply that the T4 spinal nerve dorsal ramus could be the only one that carries motor axons beyond the para-/inter-spinous muscles in addition to the usual sensory and sympathetic axons. Alternatively, if the motor innervation of the Cutaneus scapularis muscle arises from the ventral ramus of the spinal nerve then it would suggest that ventral ramus axons reform with dorsal ramus axons to some extent. Regardless, the course or pattern of innervation is likely to be unique.

### Vascular and neural supply

No obvious vascular supply was revealed from the gross dissections, but standard anatomical principles suggest that the vascular supply travels with the innervation. Based on the limited gross anatomical observations and the microscopic fluorescence imaging of Thy1-GFP and Thy1-YFP in the Cutaneous Scapularis muscle of rats and mice (respectively), neural supply likely is derived from a single nerve contacting the muscle near the middle of the belly, possibly the single structure identified in Figure 6. It is not yet clear if this structure is a nerve, and if it contains vasculature. Presuming it is indeed a nerve, it is not clear if the nerve is a branch of the T4 dorsal cutaneous nerve or a separate nerve that was closely apposed to, or even fasciculated with, the T4 DCn. Identifying the peripheral course of innervation and the location of sensory and motor axons is a priority for thorough characterization of the muscle and understanding its developmental origins, its function, and its potential utility in experimental systems.

### Function

An excellent and encompassing discussion of demonstrated and proposed functions of the cutaneous muscles has already been published so this discussion will remain more focussed [6].

On first consideration, the muscle would seem to nicely fill a gap for defensive skin movement. The Cutaneous trunci muscle is well described as providing reflexive defensive movement of skin to rid the organism of pests or other skin-contacting objects. But the CTM reflex (CTMR) does not respond to stimuli from, nor does it move, much of the cervical and upper thoracic skin [11]. The CutScap may provide a means for moving the back skin between the base of the skull and the CTm, which otherwise would have little reflexive protection.

Although this is an appealing possibility, the insertion into dermis is quite small, while the area to be moved is comparatively much larger and some of the loosest skin on the body, suggesting that such an action may not be highly effective.

We previously described the range of segmental inputs driving the CTMR and the skin areas that are moved [11]. If the Cutaneus scapularis contributes to skin movement similarly to the Cutaneus trunci, and the neuro-muscular circuitry is organized similarly and similarly resistant to pentobarbital, it stands to reason that the cutaneous nerves serving the skin near the dermal insertion would provide reflex-driving sensory input. In the pentobarbital-anesthetized rat, reflexive behavior was observed in response to pinch of the peri-midline skin extending as far rostral as the lower cervical spinal segments. The CTMR in response to natural stimulation of that lower cervical skin was restricted to the region of the CTm that was not attached to skin. Electrical stimulation of the cervical DCnn did not drive a functional CTMR, but induced weak-but-EMG-recordable contraction of the belly of the CTm which is not attached to the dermis. Those not-reduced preparations could not discriminate skin movement due to CTm and Cutaneus scapularis, but the location of skin movement suggests the Cutaneus scapularis was either not contracting, or not contracting effectively. This further suggests that stimulation of the cervical dorsal cutaneous nerves does not drive activation of the Cutaneus scapularis muscle, leaving the possibility that activation of the Cutaneus scapularis sufficient to move skin is driven by other nerves, possibly the thoracic dorsal cutaneous nerves. If this were true then Cutaneus scapularis-attached skin should have moved in the non-reduced preparations, which we did not observe. Our observations cannot address the possibility that the Cutaneus scapularis was activated in those conditions by input from the cervical or thoracic cutaneous nerves, but were insufficient to induce skin movement.

Together these observations suggest that the Cutaneous scapularis muscle may not be involved in this sort of function, or that it may not be as effective as the CTM reflex. Nonetheless, the length of the Cutaneous scapularis muscle might provide for extensive contraction and sufficient movement of very loose skin to dislodge pests, but this has yet to be determined.

It is possible that the adult version of the Cutaneus scapularis may be the residual of a muscle with a neonate-specific function. For example, the orientation suggests the possibility that the muscle may act as a suspension system for when pups are carried by the nape of their neck by their parent. Contrary to this possibility, rodents carry their pups more by the trunk than the neck. If this were the case, one would question why other species do not have this muscle despite carrying their young by the nape of the neck. One possibility is that other muscles not present in, or less-developed in, the rodents perform the same task. For example, the large platysma in the cat – a clear nape-carrier species – may serve this purpose. Another possibility is that the muscle is present in many species but recedes throughout postnatal development in non-rodents but remains in the rodents. Alternatively, the muscle may be entirely vestigial in rodents and absent in higher species.

It is also possible that the muscle functions predominantly as a sensor, similar to what has been proposed for the sternalis and serratus posterior muscles [16, 17], perhaps more likely in the neonatal than adult context.

Although more investigation is required to confirm this, from observation and manipulation it appears that contraction of the Cutaneus scapularis in the adult might impinge on the rostro-medial border of Cutaneus trunci muscle when both are contracting, which is not a common feature across any muscle type. If impingement is confirmed, this might suggest that the main purpose of the muscle is for some pre-adult function. Developmental organization may have dictated that the closest origin-point was distal to (or shared with) the Spinotrapezius, therefore committing the muscle to a significant length and unusual course in adulthood.

### Utility

At very least, a minimal characterization will provide the field with means to reduce variation and unexplained differences in outcomes that could result from the presence of an unknown muscle and innervation in supposedly fully-defined tissues. Clinically, the CTMR is often used to help define the level of spinal injuries (e.g., [18, 19]). If the Cutaneus scapularis is involved in the movement of skin in response to localized stimulation similarly to, or in conjunction with, the CTRM then the neuro-motor circuitry of that response must be defined in order to not mislead diagnoses.

Regarding more basic science, for example, single-cell RNA-sequencing is starting to define motor neuron types with increasing specificity, in some cases identifying new cell types [20-25]. The power of these definitions for peripheral nervous system neurons relies on an understanding of the targets of the neurons. Certainly an RNA-seq-identified novel class of neuron can lead to seeking novel or more specific target information. However, a greater *a priori* understanding of which targets are innervated by the motoneurons in each segment could aid interpretation. In addition, the intrinsic regenerative capabilities and mechanisms of cutaneous muscles differ in useful ways [26], enhancing the importance of clearly defining the range of cutaneous muscles.

If initial observations are borne out, this novel muscle could represent a unique system in many ways. The T4 spinal nerve dorsal ramus could be the only one that carries motor axons beyond the epaxial paraspinal muscles in addition to the usual sensory and sympathetic axons [27-29], offering a comparative model in the spinal system to identify factors controlling axon sorting and pathfinding during development. The Cutaneus scapularis already appears to represent the only dorsal cutaneous muscle with a caudal-to-rostral orientation. These features again could offer a comparative model system for understanding developmental controls. Study of cranial cutaneous musculature is important for a variety of pathologies and therapeutic development, but is notoriously difficult because of the small size of most animal models. The more accessible cutaneous muscles and easy-to-label/obtain motoneurons of the trunk cutaneous muscles may offer an attractive complementary system.

Far more length of the Cutaneus scapularis is easily accessed whereas the vast majority of other cutaneous muscles, including large muscles such as the CTm, are adhered to dermis. The Cutaneus scapularis can be easily removed for a variety of purposes (ex vivo physiology, transplantation, etc.) with very little impact on the functioning of the animal, unlike muscles of similar length (e.g., sartorius) which have ligamentous connection to bone.

### Community Engagement

These findings were prepared and placed on BioRXiv to facilitate community engagement while further characterization is undertaken. Anyone with knowledge of prior descriptions of this muscle in rodents or other species, or homologs and analogs in other species, is encouraged to provide comments or contact the author.

## Acknowledgements

The author thanks anatomical expert colleagues Nicole Herring and Jennifer Brueckner-Collins at the University of Louisville, Richard D. Johnson at the University of Florida, and attendees of the 2019 joint meeting of the Human Anatomy and Physiology Society (HAPS) and the American Association of Clinical Anatomists (AACA) at Bellarmine University in Louisville, KY for their consideration of the identity of the muscle and possible homologues or analogues. Thanks also to Drs. Dena Howland and Teresa Pitts of the University of Louisville for allowing dissection of post-mortem cats used in separate experiments. Thanks also to Dr. Matt Wood and J. Blake Schoenfeld of Washington University in St. Louis for use of Thy1-GFP rat tissue.

## Funding

This work was supported in part by NIH awards R01NS094741 and R21120498 and VA I21RX003766

## Disclosures

The author is a founder of dSentz, Inc. and Pioneer Neurotech, Inc. The work reported herein is entirely separate from either company. Neither company exerted any editorial oversight. There are no conflicts of interest to disclose.

## Notes

### Competing Interest Statement

The authors have declared no competing interest.

